# Synthetic Lethality of Wnt Pathway Activation and Asparaginase in Drug-Resistant Acute Leukemias

**DOI:** 10.1101/415711

**Authors:** Laura Hinze, Maren Pfirrmann, Salmaan Karim, James Degar, Connor McGuckin, Divya Vinjamur, Joshua Sacher, Kristen E. Stevenson, Donna S. Neuberg, Daniel E. Bauer, Florence Wagner, Kimberly Stegmaier, Alejandro Gutierrez

## Abstract

Resistance to asparaginase, an antileukemic enzyme that depletes asparagine, is a common clinical problem. Using a genome-wide CRISPR/Cas9 screen, we found a synthetic lethal interaction between Wnt pathway activation and asparaginase in acute leukemias resistant to this enzyme. Wnt pathway activation induced asparaginase sensitivity in distinct treatment-resistant subtypes of acute leukemia, including T-lymphoblastic, hypodiploid B-lymphoblastic, and acute myeloid leukemias, but not in normal hematopoietic progenitors. Sensitization to asparaginase was mediated by Wnt-dependent stabilization of proteins (Wnt/STOP), which inhibits GSK3-dependent protein ubiquitination and degradation. Inhibiting the alpha isoform of GSK3 phenocopied this effect, and pharmacologic GSK3α inhibition profoundly sensitized drug-resistant leukemias to asparaginase. Our findings provide a molecular rationale for activation of Wnt/STOP signaling to improve the therapeutic index of asparaginase.

**SIGNIFICANCE:** The intensification of asparaginase-based therapy has improved outcomes for several subtypes of acute leukemia, but the development of treatment resistance has a poor prognosis. We hypothesized, from the concept of synthetic lethality, that gain-of-fitness alterations in drug-resistant cells had conferred a survival advantage that could be exploited therapeutically. We found a synthetic lethal interaction between activation of Wnt-dependent stabilization of proteins (Wnt/STOP) and asparaginase in acute leukemias resistant to this enzyme. Inhibition of the alpha isoform of GSK3 was sufficient to phenocopy this effect, and the combination of GSK3α-selective inhibitors and asparaginase had marked therapeutic activity against leukemias resistant to monotherapy with either agent. These data indicate that drug-drug synthetic lethal interactions can improve the therapeutic index of cancer therapy.

## INTRODUCTION

The genetic concept of synthetic lethality describes an interaction between two mutations that are each well-tolerated individually, but are lethal when combined [reviewed in (Ashworth and Lord, 2018)]. Small molecule inhibitors can phenocopy the effect of specific mutations, which led to the search for synthetic lethal drug-mutation interactions that would be selectively toxic to cancer cells harboring specific oncogenic mutations (Hartwell et al., 1997). Proof-of-principle for this idea came from the discovery that PARP inhibitors are profoundly toxic to tumor cells harboring biallelic BRCA1/2 gene mutations, but not to normal cells that retain at least one functional allele, thus providing a striking therapeutic index (Farmer et al., 2005). Inspired by this concept, we reasoned that drug-drug synthetic lethal interactions had the potential to improve the therapeutic index of cancer therapy, if applied to drugs that were sufficiently selective for cancer cells.

Asparaginase, an exogenous enzyme that deaminates the nonessential amino acid asparagine, has long been recognized to have activity against aggressive hematopoietic neoplasms (Broome, 1961). Asparaginase dose-intensification has improved outcomes for T-cell and B-cell acute lymphoblastic leukemias (T-ALL and B-ALL) (Clavell et al., 1986; DeAngelo et al., 2015; Ertel et al., 1979; Pession et al., 2005). This enzyme also has therapeutic activity in acute myeloid leukemias and in some non-Hodgkin lymphomas (Capizzi et al., 1988; Wells et al., 1993; Yamaguchi et al., 2011). The development of resistance to asparaginase-based treatment regimens has a poor prognosis, and effective therapeutic options are lacking for many of these patients.

The sensitivity of acute leukemia cells to asparaginase is due, at least in part, to low expression of asparagine synthetase (ASNS) by these cells, resulting in their dependence on exogenous asparagine (Haskell and Canellos, 1969; Horowitz et al., 1968). By contrast, physiologic expression of *ASNS* by most normal cells is thought to explain the favorable therapeutic index of asparaginase (Rizzari et al., 2013). Increased ASNS expression by leukemic blasts can induce asparaginase resistance (Haskell and Canellos, 1969; Horowitz et al., 1968). However, ASNS is not an ideal therapeutic target because its inhibition is likely to worsen asparaginase-induced toxicity to normal tissues. Nevertheless, *ASNS* expression is not the sole determinant of asparaginase response (Appel et al., 2006; Hermanova et al., 2012; Holleman et al., 2004; Stams et al., 2003), whose biologic basis remains incompletely understood.

We hypothesized that asparaginase-resistant leukemia cells harbor gain-of-fitness alterations whose therapeutic targeting would be uniquely toxic to tumor cells upon treatment with the enzyme. Thus, we performed a genome-wide loss-of-function genetic screen in asparaginase-resistant T-ALL cells. This revealed that negative regulators of Wnt signaling were selectively depleted upon treatment with asparaginase. Wnt pathway activation induced asparaginase sensitization in distinct treatment-resistant subtypes of acute leukemia, including T-ALL, hypodiploid B-ALL and acute myeloid leukemia. This effect was independent of canonical Wnt/β-catenin or mTOR activation. Instead, it was mediated by Wnt-dependent stabilization of proteins (Wnt/STOP), which inhibits GSK3-dependent protein ubiquitination and degradation. Inhibition of the alpha isoform of GSK3 was sufficient to phenocopy this effect, and the combination of GSK3α inhibitors with asparaginase had profound therapeutic activity against leukemias that were completely refractory to monotherapy with either of these drugs. Together, these results demonstrate a synthetic lethal interaction between asparaginase and activation of Wnt/STOP signaling.

## RESULTS

### Wnt Pathway Activation Induces Asparaginase Sensitization

To identify molecular pathways that promote leukemic cell fitness upon treatment with asparaginase, we performed a genome-wide CRISPR/Cas9 loss-of-function genetic screen in the T-ALL cell line CCRF-CEM, which is asparaginase-resistant (Figure S1). We first optimized conditions for a drop-out screen using positive control guide RNAs targeting asparagine synthetase (*ASNS*), the enzyme that synthesizes asparagine (Figure S2A-C) (Van Heeke and Schuster, 1989). We then transduced Cas9-expressing CCRF-CEM cells with the GeCKO genome-wide guide RNA library (Shalem et al., 2014) (Figure 1A, Figure S2D), treated with either vehicle or a 10 U/L dose of asparaginase that lacked detectable toxicity, and guide RNA representation was assessed. *ASNS* was the gene most significantly depleted in asparaginase-treated cells, followed closely by two regulators of Wnt signaling, *NKD2* and *LGR6* (Figure 1B). NKD2 negatively regulates Wnt signaling by binding and repressing dishevelled proteins (Wharton et al., 2001), whereas LGR proteins function as R-spondin receptors reported to either activate or repress Wnt signaling (de Lau et al., 2011; Walker et al., 2011). To test how these genes regulate Wnt signaling, we transduced CCRF-CEM cells with shRNAs targeting *NKD2* or *LGR6*, or with an shLuciferase control (Figure 1C). Knockdown of *NKD2* or *LGR6* increased levels of active (nonphosphorylated) β-catenin (Figure 1D), as well as the activity of a TOPFlash reporter of canonical Wnt/β-catenin transcriptional activity (Fuerer and Nusse, 2010) (Figure 1E). Thus, *NKD2* and *LGR6* are negative regulators of Wnt signaling in T-ALL cells. To validate that loss of *NKD2* or *LGR6* sensitizes these cells to asparaginase, we again transduced CCRF-CEM cells with these shRNAs and treated them with asparaginase. Knockdown of *NDK2* or *LGR6* profoundly sensitized the cells to asparaginase (Figure 1F), and potentiated asparaginase-induced caspase activation, indicating induction of apoptosis (Figure 1G). This effect was phenocopied by treatment with the Wnt ligand Wnt3A (Figure 1H). Thus, Wnt pathway activation sensitizes T-ALL cells to asparaginase.

**Figure 1.**
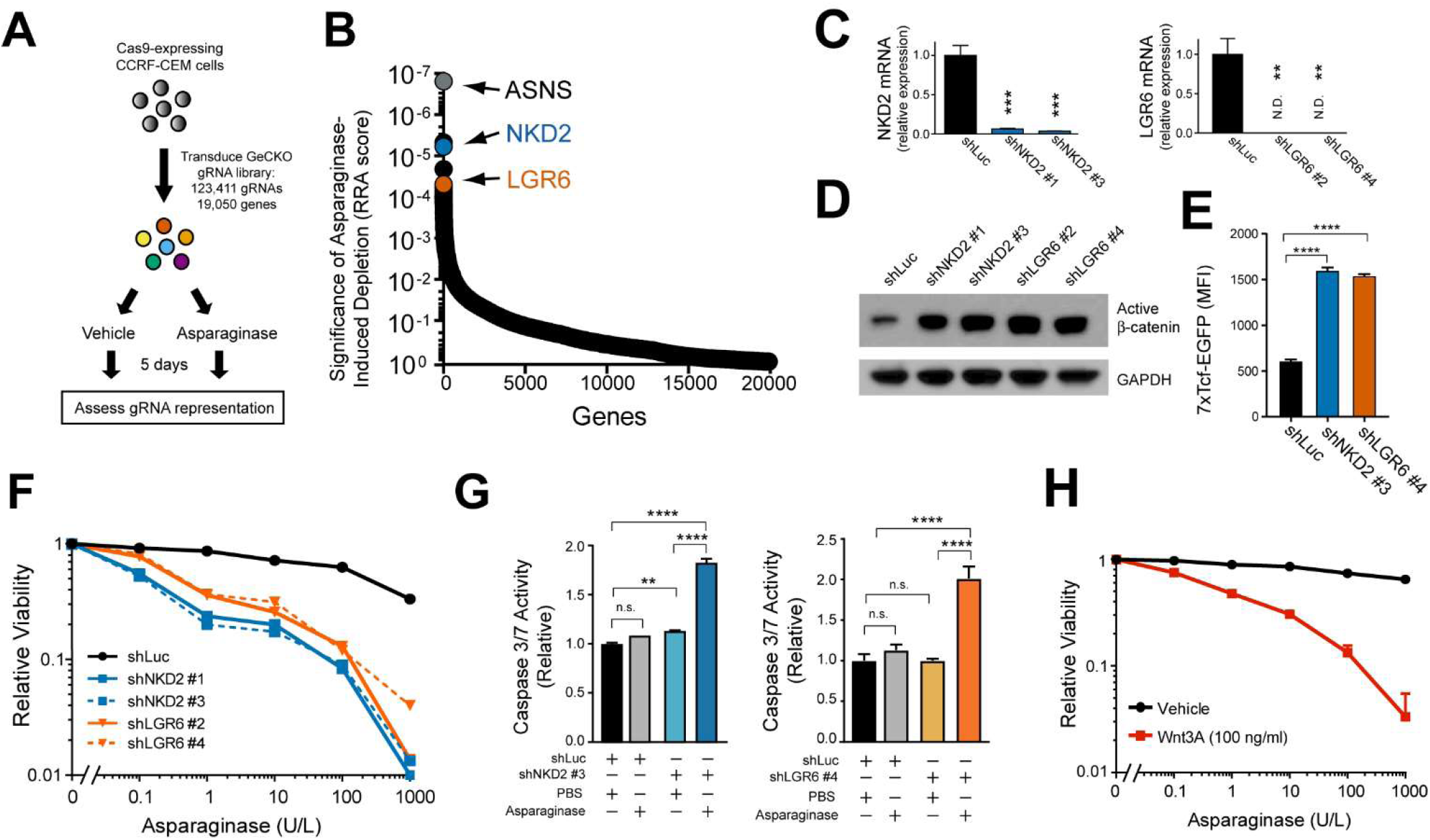
Wnt Pathway Activation Sensitizes Leukemia Cells to Asparaginase. (A) CCRF-CEM cells were transduced with the GeCKO genome-wide guide RNA library in biologic duplicates, split into treatment with vehicle or asparaginase (10 U/L), and guide RNA representation was assessed after 5 days of treatment. (B) Significance of gene depletion in asparaginase-treated conditions, as assessed by robust ranking aggregation (RRA) score calculated using MAGeCK analysis. Note that microRNA genes are not shown. (C) CCRF-CEM cells were transduced with the indicated shRNAs, and knockdown efficiency was assessed by RT-PCR analysis. CT values greater than 36 were defined as not detected (N.D.). (D) CCRF-CEM cells were transduced with the indicated shRNAs, and subjected to Western blot analysis for active (nonphosphorylated) β-catenin or GAPDH. (E) CCRF-CEM cells expressing a lentiviral 7xTcf-EGFP (TopFLASH) reporter were transduced with the indicated shRNAs, and reporter-driven EGFP fluorescence was assessed. (F) CCRF-CEM cells were transduced with the indicated shRNAs and treated with the indicated doses of asparaginase. Relative viability was assessed after 8 days of treatment by counting viable cells. All cell counts were normalized to those in shLuc-transduced, no-asparaginase controls. (G) CCRF-CEM cells were transduced with the indicated shRNAs, treated with asparaginase (10 U/L) for 48 hours, and caspase 3/7 activity assay was assessed. (H) CCRF-CEM cells were treated as indicated and the number of viable cells after 8 days of treatment was assessed. All cell counts were normalized to those in no-Wnt3a, no-asparaginase controls.

### GSK3 Inhibition Sensitizes Distinct Acute Leukemia Subtypes to Asparaginase-Induced Cytotoxicity

Inhibition of glycogen synthase kinase 3 (GSK3) activity is a key event in Wnt-induced signal transduction (Nusse and Clevers, 2017; Siegfried et al., 1992; Taelman et al., 2010), prompting us to test whether this effect could be phenocopied by CHIR99021, an ATP-competitive inhibitor of both GSK3 paralogs, GSK3α and GSK3β(Bennett et al., 2002). CHIR99021 induced significant asparaginase sensitization across a panel of cell lines representing distinct subtypes of treatment-resistant acute leukemia, including T-ALL, acute myeloid leukemia and hypodiploid B-ALL (Figure 2A and 2B). Importantly, CHIR99021 did not increase the toxicity of asparaginase to normal human CD34+ hematopoietic progenitors (Figure 2C), suggesting a selective effect on leukemic cells. Nor did it sensitize leukemic cells to other commonly used antileukemic drugs, including vincristine, 6-mercaptopurine, dexamethasone, and doxorubicin (Figure 2D). Thus, GSK3 inhibition can enhance asparaginase toxicity not only in T-ALL, but in other common acute leukemia variants as well.

**Figure 2.**
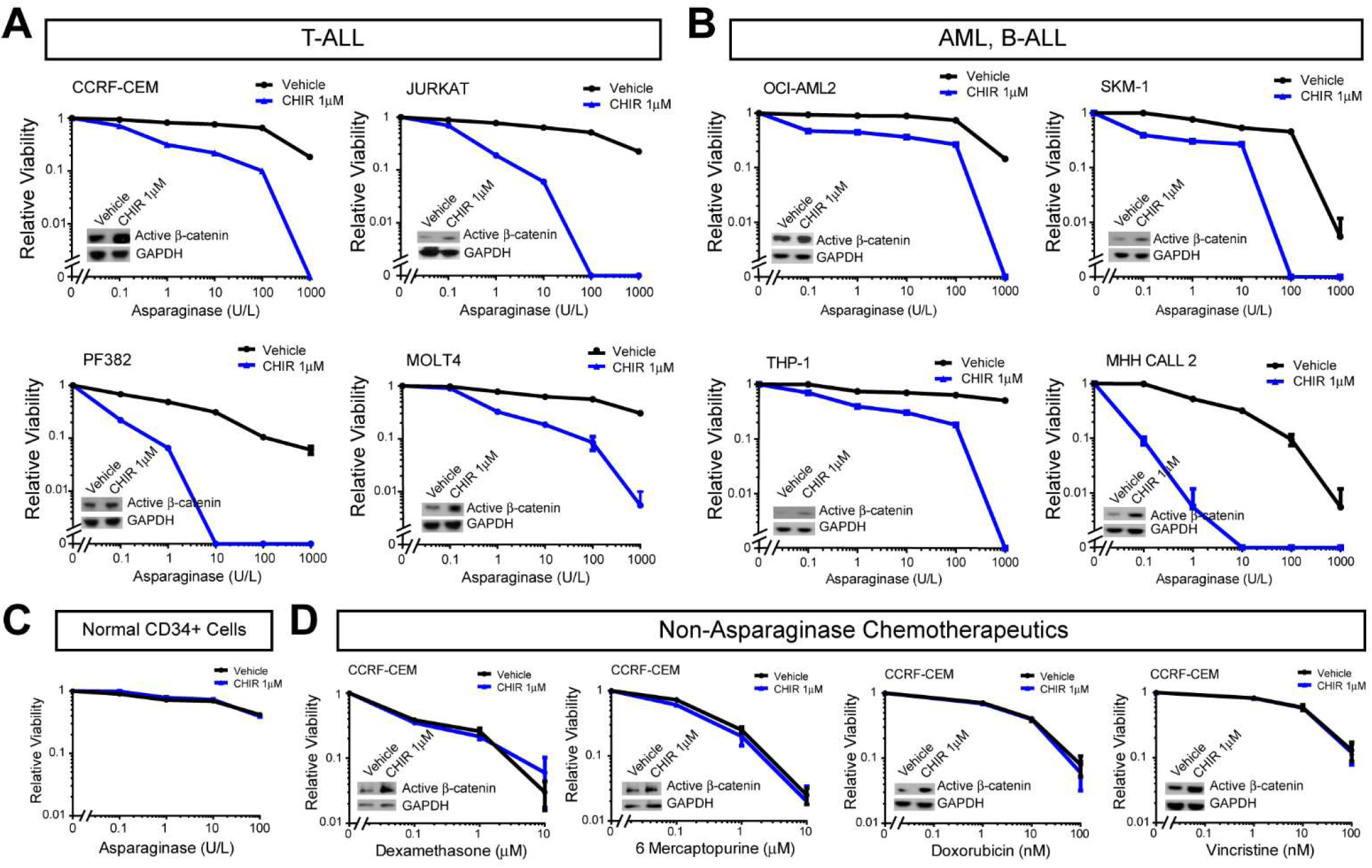
GSK3 Inhibition Sensitizes Distinct Acute Leukemia Subtypes, but not Normal Hematopoietic Progenitors, to Asparaginase-Induced Cytotoxicity. (A-B) The indicated cells were treated with the GSK3 inhibitor CHIR99021 (CHIR, 1 μM) or vehicle control, together with the indicated doses of asparaginase for 8 days. Relative viability was assessed based on viable cell counts, all of which were normalized to those in vehicle-treated cells. (C) Normal CD34+ human hematopoietic progenitor cells were treated with CHIR99021 (1 μM) or vehicle, together with the indicated doses of asparaginase, and viable cell counts were assessed after treatment for 4 days. Note that these normal hematopoietic progenitors could not be maintained for more than 4 days in culture. (D) CCRF-CEM cells were treated with CHIR99021 (1 μM) or vehicle, and the indicated chemotherapeutic drugs for 8 days. All Western Blots shown indicate levels of active (nonphosphorylated) β catenin or GAPDH after treatment with CHIR 99021 (1 µM) or vehicle. Viable cells were counted and results normalized to counts in no-CHIR, no-chemotherapy controls.

### Wnt-Dependent Stabilization of Proteins Mediates Sensitization to Asparaginase

Canonical Wnt signaling is best known as an activator of β-catenin-dependent transcriptional activity (Brunner et al., 1997; van de Wetering et al., 1997), leading us to ask whether β-catenin activation is sufficient to induce asparaginase sensitization. Thus, we transduced CCRF-CEM cells with a constitutively active ΔN90 β-catenin allele (Guo et al., 2012), or shRNA targeting the β catenin antagonist gene *APC* (Moon and Miller, 1997), and treated them with asparaginase. Surprisingly, these modifications lacked any discernible effect on asparaginase sensitivity, despite effective activation of β-catenin-induced transcription, as assessed by TOPFlash reporter activity (Figure 3A and 3B). The Wnt pathway also regulates protein synthesis and metabolism by activating both mTOR complexes, mTORC1 (Inoki et al., 2006) and mTORC2 (Esen et al., 2013). However, treatment with the mTORC1 inhibitors rapamycin and RAD001, or with the dual mTORC1/2 inhibitor AZD2014, had no effect Wnt-induced sensitization to asparaginase (Figure 3C and Figure S3). On balance, these data indicate that asparaginase sensitization is independent of β-catenin or mTOR activity.

Activation of Wnt signaling increases total cellular protein content and cell size by inhibiting GSK3-dependent protein ubiquitination and proteasomal degradation, an effect termed Wnt-dependent stabilization of proteins (Wnt/STOP) (Acebron et al., 2014; Huang et al., 2015; Taelman et al., 2010). This function of the Wnt pathway appeared relevant to asparaginase sensitization, because we observed a measurable decrease in cell size after treatment with the enzyme, even in CCRF-CEM cells that are resistant to asparaginase-induced cytotoxicity (Figure 3D), and this effect was reversed by Wnt pathway activation (Figure 3E). To test whether Wnt pathway activation inhibits protein degradation in asparaginase-treated cells, we used a pulse-chase experiment to measure total protein half-life. CCRF-CEM cells were first transduced with Wnt-activating shRNAs targeting *NKD2* and *LGR6*, or with an shLuciferase control. Cells were then incubated with a pulse of the methionine analog azidohomoalanine (AHA), followed by a chase in which the cells were treated with asparaginase, and the rate of AHA release due to protein degradation was measured by flow cytometry. Wnt pathway activation did not affect the degree of AHA label incorporation during the pulse period (Figure S4); however, shRNA knockdown of *LGR6* or *NDK2* increased total protein half-life by approximately 40% in these cells (Figure 3F). These findings indicate that Wnt pathway activation inhibits protein degradation in asparaginase-treated cells.

**Figure 3.**
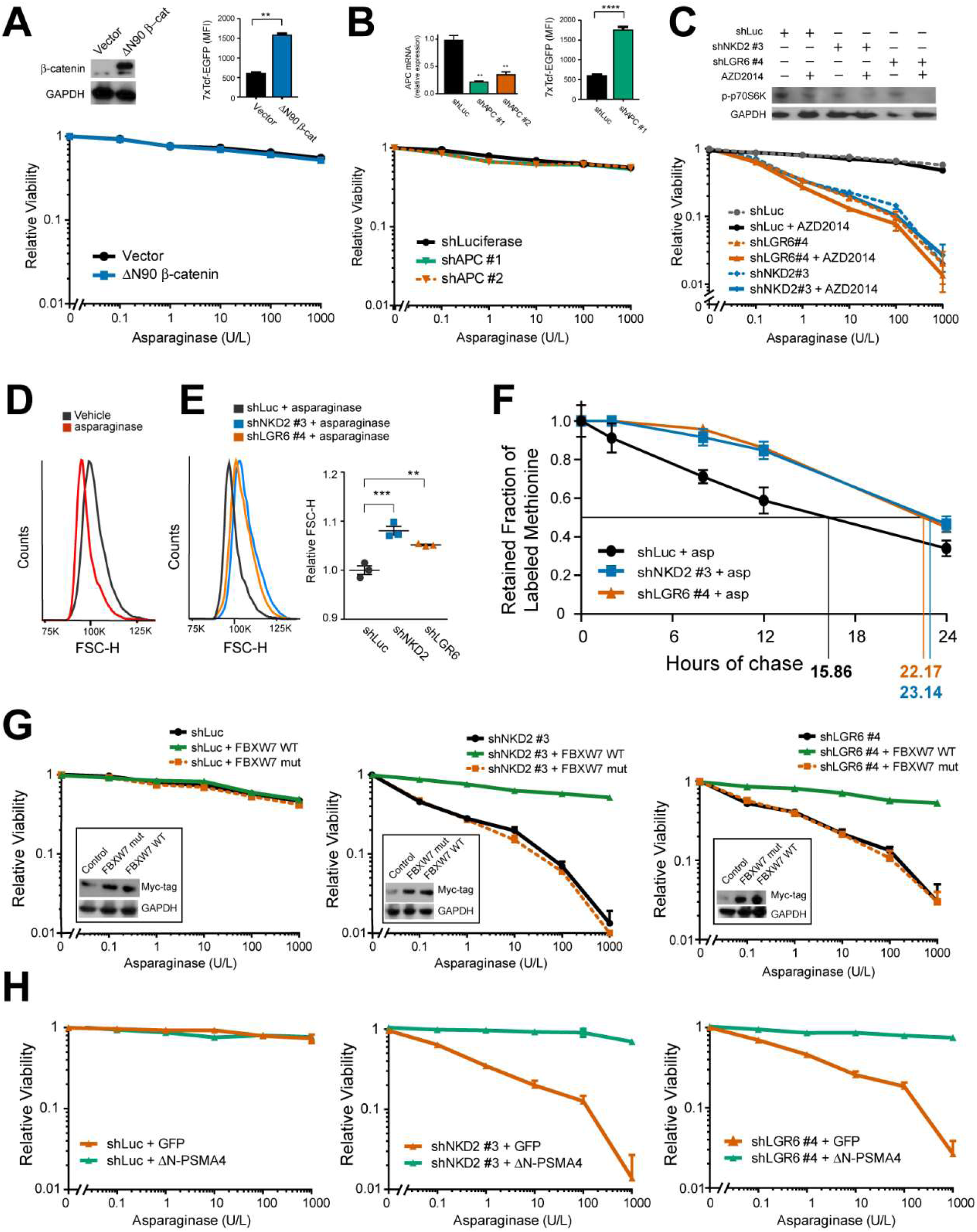
Wnt-Dependent Stabilization of Proteins Mediates Sensitization to Asparaginase. (A-B) CCRF-CEM cells transduced with the indicated constructs were analyzed by Western blot or RT-PCR, and effects on 7xTcf-EGFP Wnt reporter activity were assessed (top). Cells were treated with the indicated doses of asparaginase for 8 days, and the number of viable cells was counted. All cell counts were normalized to those in control-transduced, no-asparaginase cells. (A) CCRF-CEM cells transduced with the indicated shRNAs were treated with vehicle or AZD2014 (10 nM), and analyzed by Western blot (top). Cells were treated with the indicated doses of asparaginase and viability was assessed as in (A). (D) Cell size was assessed in CCRF-CEM cells treated with vehicle or asparaginase (10 U/L) for 48 hours by flow cytometry. (E) CCRF-CEM cells transduced with the indicated shRNAs were treated and analyzed as in (D). (F) CCRF-CEM cells transduced with the indicated shRNAs were incubated with a pulse of the methionine analog AHA, then released from AHA and treated with asparaginase (10 U/L) during the chase period. The degree of AHA label retention was assessed by flow cytometry. (G) CCRF-CEM cells transduced with the indicated constructs were analyzed by Western blot, treated with the indicated doses of asparaginase, and viability was assessed as in (A). Note that wild-type and mutant FBXW7 were Myc-tagged. (H) CCRF-CEM cells transduced with the indicated constructs were treated with the indicated doses of asparaginase, and viability was assessed as in (A).

Does inhibition of protein degradation in leukemic cells mediate sensitization to asparaginase? The E3 ubiquitin ligase component FBXW7 recognizes a canonical phospho-degron that is phosphorylated by GSK3, and overexpression of FBXW7 restores the degradation of a subset of proteins stabilized by Wnt/STOP signaling (Acebron et al., 2014). We thus asked whether FBXW7 overexpression might also reverse Wnt/STOP induced asparaginase sensitization. After transducing CCRF-CEM with Wnt-activating shRNAs or an shLuciferase control, we transduced the same cells with expression constructs encoding either wild-type FBXW7, or an FBXW7 R465C point mutant allele with impaired binding to its canonical phospho-degron (Koepp et al., 2001). Overexpression of wild-type FBXW7 blocked the ability of Wnt pathway activation to trigger asparaginase hypersensitivity, whereas the R465C mutant had no effect (Figure 3G). However, FBXW7 is a T-ALL tumor suppressor that can target specific oncoproteins (Davis et al., 2014; Thompson et al., 2007), prompting us to test whether direct stimulation of proteasomal activity is sufficient to reverse Wnt-induced sensitization to asparaginase. Thus, we leveraged a hyperactive open-gate mutant of the proteasomal subunit PSMA4 (ΔN-PSMA4), whose expression is sufficient to stimulate degradation of a wide range of proteasomal substrates (Choi et al., 2016). CCRF-CEM cells were first transduced with control or Wnt-activating shRNAs, and then transduced with the constitutively active proteasomal subunit ΔN-PSMA4, or a GFP control. Expression of the constitutively active ΔN-PSMA4 proteasomal subunit completely blocked Wnt-induced sensitization to asparaginase (Figure 3H). These findings indicate that Wnt pathway activation sensitizes leukemic cells to asparaginase by inhibiting proteasomal degradation of proteins.

### Synthetic Lethality of GSK3α Inhibition and Asparaginase in Human Leukemia

To explore the therapeutic potential of our studies, we focused on GSK3 as a therapeutic target. Although the pan-GSK3 inhibitor that we used in cell lines lacks suitable pharmacokinetic properties for in vivo studies, isoform-selective inhibitors of each GSK3 paralog with favorable pharmacologic properties have recently been developed (Wagner et al., 2018). We found that the GSK3α-selective inhibitor BRD0705 effectively sensitized CCRF-CEM cells to asparaginase, while the GSK3 β-selective inhibitor BRD3731 had only modest effects (Figure 4A). To determine whether this result reflects redundancy between GSK3 isoforms, or partial inhibition of GSK3α by the GSK3 β inhibitor BRD3731, we used shRNAs to selectively deplete GSK3α or GSK3 β. Depletion of GSK3α phenocopied Wnt-induced sensitization to asparaginase, whereas GSK3 β knockdown had no effect (Figure 4B). Importantly, the effect of GSK3α-targeting shRNAs was rescued by transduction of a GSK3α expression construct that escapes shRNA targeting (Figure 4C).

**Figure 4.**
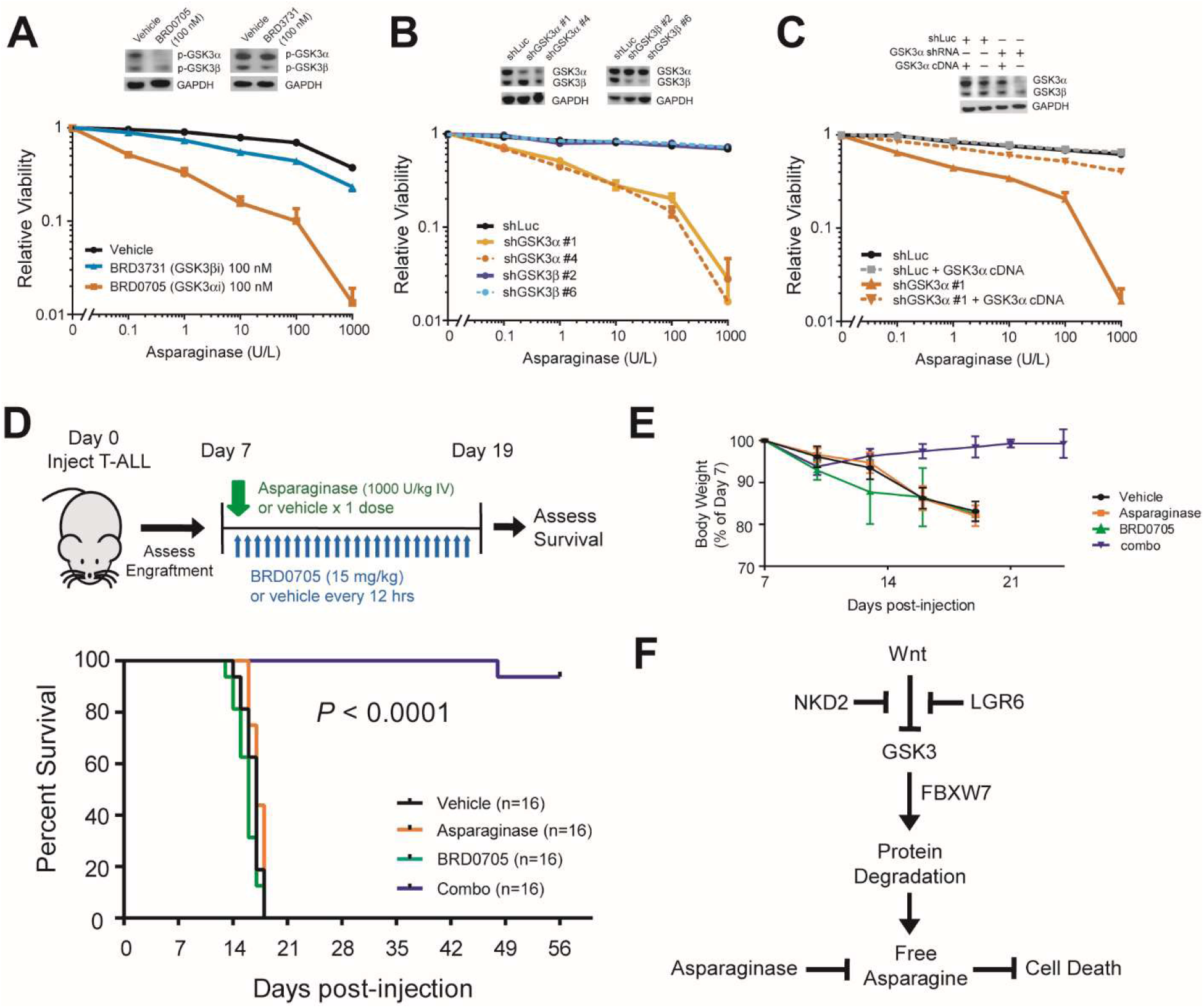
Synthetic Lethality of GSK3α Inhibition and Asparaginase in Human Leukemia. (A) CCRF-CEM cells were treated with vehicle, the GSK3α-selective inhibitor BRD0705, or the GSK3α-selective inhibitor BRD3731, and analyzed by Western blot analysis (inset) for phospho-GSK3 or GAPDH. Cells were then treated with the indicated drugs 8 days, and the number of viable cells was counted. All cell counts were normalized to those in vehicle-treated controls. (B) CCRF-CEM cells were transduced with the indicated shRNAs, and analyzed by Western blot (inset) for total GSK3 or GAPDH. Cells were then treated with the indicated doses of asparaginase, and viability was assessed as in (A). (C) CCRF-CEM cells were transduced with the indicated shRNAs, without or with a GSK3α expression construct that escapes shRNA targeting. Western blot analysis (inset) was performed for total GSK3 or GAPDH. Cells were then treated with the indicated doses of asparaginase, and viability was assessed as in (A). (D) T-ALL cells from a patient-derived xenograft were injected into NRG immunodeficient mice. After engraftment of leukemia (>5% human leukemic cells in peripheral blood), mice were treated as indicated. Differences in survival of mice treated with vehicle, asparaginase alone, or BRD0705 alone were not significant. Differences between each of these groups and mice treated with the combination of asparaginase and BRD0705 (combo) were all P < 0.0001. (E) Weights of mice in the experiment shown in (D). Note that treatment started on day 7. (F) Proposed model.

Combined treatment with the GSK3α inhibitor BRD0705 and asparaginase was well-tolerated by NRG mice, with no appreciable weight changes or increases in serum bilirubin levels, an important dose-limiting toxicity of asparaginase in adults (data not shown). Finally, we injected a cohort of NRG mice with leukemic cells from a primary asparaginase-resistant T-ALL patient-derived xenograft. Once leukemia engraftment was detected in the peripheral blood, we treated the mice with vehicle, asparaginase, BRD0705, or the combination of asparaginase and BRD0705. Although the xenografts proved highly resistant to asparaginase or BRD0705 monotherapy, the combination of asparaginase and BRD0705 was both highly efficacious and well-tolerated (Figure 4D and 4E).

## DISCUSSION

Using a genome-wide CRISPR/Cas9 screen, we identified a synthetic lethal interaction between activation of Wnt signaling and asparaginase in acute leukemias resistant to this enzyme. Wnt activation results in inhibition of GSK3 activity (Siegfried et al., 1992; Taelman et al., 2010), and pharmacologic GSK3 inhibition sensitized a panel of treatment resistant acute leukemias to asparaginase. The effect of Wnt signaling was independent of β-catenin or mTOR activation. Instead, we found that activation of Wnt/STOP signaling mediates sensitization to asparaginase. Indeed, expression of a hyperactive proteasomal subunit that broadly increases proteasomal degradation (Choi et al., 2016) completely blocked Wnt-induced sensitization to asparaginase. These findings support a model in which regulated protein degradation is an important source of asparagine in drug-resistant leukemias treated with asparaginase, an adaptive pathway that is blocked by activation of Wnt/STOP signaling (Figure 4F).

These results demonstrate that synthetic lethal drug-drug interactions can be leveraged to improve the therapeutic index of cancer therapy. This approach requires drug combinations with selective toxicity to cancer cells. We anticipate several classes of therapeutics that may be particularly well-suited for such an approach: drugs devised from the principles of drug-mutation synthetic lethal interactions, such as PARP inhibitors for BRCA mutant tumors (Farmer et al., 2005); drugs that target mutant oncoproteins that are uniquely present in cancer cells, such as selective inhibitors of neomorphic IDH gene mutations (Wang et al., 2013); or drugs that target genetic vulnerabilities induced by “passenger” gene mutations, such as those targeting MTAP deletions acquired due to its proximity to the CDKN2A tumor suppressor (Kryukov et al., 2016). It is perhaps less predictable that such an approach would be successful for conventional chemotherapeutics, which generally target cellular factors that are required for both normal and malignant cells. Nevertheless, the fact that these drugs are the primary therapeutic option with curative potential for a broad range of human cancers demonstrates the existence of a substantial therapeutic window, which can be further improved as demonstrated here.

It is of considerable interest that the combination of Wnt/STOP activation and asparaginase is potently toxic to leukemic cells, whereas normal cells appear to have effective compensatory mechanisms. One possibility is that normal cells have an improved capacity for asparagine biosynthesis, and are thus not dependent on regulated protein degradation to tolerate asparaginase. However, several of the asparaginase-resistant cells utilized in our studies express asparagine synthetase, suggesting a role for additional factors. These may include checkpoints that normally couple proliferation to asparagine availability, a link that could be readily disrupted by oncogenic mutations. If so, the combination of Wnt/STOP activation and asparaginase could have efficacy against a broad range of tumors harboring relevant gene mutations.

## SUPPLEMENTAL INFORMATION

Supplemental Information includes four supplemental figures and one supplemental table (Table S1, Significance of gene depletion or enrichment in asparaginase-treated cells. Related to Figure 1) and can be found with this article online.

## ACKNOWLEDGEMENTS

We thank Alex Kentsis, Ralph DeBerardinis, Marc Mansour, and Min Jae Lee for advice and discussion, and Meaghan McGuinness, Natsuko Yamagata, Ronald Mathieu, Mahnaz Paktinat, Zachary Herbert and Peter Blanding for experimental assistance. We thank John Gilbert for editorial assistance. This work was supported by NIH/NCI grant 1R01CA193651 and the Boston Children’s Hospital Translational Investigator Service. L.H. was supported by the German National Academic Foundation and the Biomedical Education Program.

## AUTHOR CONTRIBUTIONS

The experiments were conceived and designed by L.H., M.P. and A.G. L.H., M.P., S.K., and J.D. performed experiments and analyzed data. L.H., M.P., D.V., C.M., J.S., K.E.S., D.S.N., D.E.B., F.W., K.S., and A.G. designed experiments and analyzed data. L.H. and A.G. wrote the manuscript with input from all authors.

## METHODS

### Cell Lines and Cell Culture

293T cells, T-ALL cell lines, AML cell lines and B-ALL cell lines were obtained from ATCC (Manassas, VA, USA), DSMZ (Braunschweig, Germany), the A. Thomas Look laboratory (Boston, MA, USA) or Alex Kentsis laboratory (New York, NY, USA) and cultured in DMEM, RPMI 1640 or MEM alpha (Thermo Fisher Scientific) with 10% or 20% fetal bovine serum (FBS, Sigma-Aldrich, Saint Louis, MO) or TET system approved FBS (Clontech, Mountain View, CA) and 1% penicillin/streptomycin (Thermo Fisher Scientific) at 37°C, 5% CO2. Human CD34+ progenitor cells from mobilized peripheral blood of healthy donors were obtained from Fred Hutchinson Cancer Research Center (Seattle, WA, USA). CD34+ progenitors were cultured in IMDM (Thermo Fisher Scientific) supplemented with 20% FBS and recombinant human interleukin-3 (R&D systems, Minneapolis, MN), recombinant human interleukin-6 (R&D systems, Minneapolis, MN) and recombinant human stem cell factor (R&D systems, Minneapolis, MN) to a final concentration of 50 ng/ml each.

Cell line identities were validated using STR profiling at the Dana-Farber Cancer Institute Molecular Diagnostics Laboratory (most recently in June 2018), and mycoplasma contamination was excluded using the MycoAlert Mycoplasma Detection Kit according to the manufacturer’s instructions (Lonza, Portsmouth, NH; most recently in March 2018).

### Mice

NOD rag gamma (NRG) mice were purchased from the Jackson Laboratories (Bar Harbor, ME; Stock # 007799). 6-8 weeks-old female NRG mice were used for experiments and littermates were kept in individual cages. Mice were randomly assigned to experimental groups. Mice were handled in strict accordance with Good Animal Practice as defined by the Office of Laboratory Animal Welfare. All animal work was done with Boston Children’s Hospital (BCH) Institutional Animal Care and Use Committee approval (protocol # 15-10-3058R).

### Lentiviral and Retroviral Transduction

Lentiviruses were generated by co-transfecting pLKO.1 plasmids of interest together with packaging vectors psPAX2 (a gift from Didier Trono; addgene plasmid # 12260) and VSV.G (a gift from Tannishtha Reya; addgene plasmid # 14888) using OptiMEM (Invitrogen, Carlsbad, CA) and Fugene (Promega, Madison, WI), as previously described (Burns et al., 2018). Retrovirus was produced by co-transfecting plasmids with packaging vectors gag/pol (a gift from Tannishtha Reya; addgene plasmid # 14887) and VSV.G.

Lentiviral and retroviral infections were performed by spinoculating T-ALL cell lines with virus-containing media (1,500 g × 90 minutes) in the presence of 8 μg/ml polybrene (Merck Millipore, Darmstadt, Germany). Selection with antibiotics was started 24 hours after infection with neomycin (700 μg/ml for a minimum of 5 days; Thermo Fisher Scientific), puromycin (1 μg/ml for a minimum of 48 hours; Thermo Fisher Scientific), or blasticidin (15 μg/ml for a minimum of 5 days; Invivogen).

Transient transfection was performed using Lipofectamine 2000 reagent (Invitrogen). Briefly, 800,000 cells were seeded in 2ml of growth medium in 24 well plates. Five μg plasmid of interest and 10 μl lipofectamine were mixed with 300 μl OptiMEM, incubated for 10 minutes and added to the wells. Antibiotic selection was begun after 48 hours of incubation.

### Pooled ASNS/AAVS1 Library

A pooled guide RNA library with 3 unique guide RNAs targeting genomic loci encoding the catalytic domain of *ASNS* (http://www.uniprot.org/uniprot/P08243), and 3 unique guide RNAs targeting the safe-harbor AAVS1 locus located in intron 1 of the *PPP1R12C* gene (Sadelain et al., 2011), were designed as described (Sanjana et al., 2014). The oligos were cloned to a modified version of lentiGuide-Puro (a gift from Feng Zhang, addgene plasmid # 52963) in which the guide RNA scaffold was replaced by a structurally optimized form (A-U flip and stem extension, called combined modification) previously reported to increase the efficiency of Cas9 targeting (Chen et al., 2013). Briefly, guide RNAs targeting the relevant genomic loci were designed using the Zhang lab CRISPR design tool (http://crispr.mit.edu/). One μl of 100 μm forward and reverse oligo was mixed with 1 μl 10X T4 DNA Ligation Buffer (NEB, Ipswich, MA),6.5 μl ddH2O, 0.5 μl T4 PNK (NEB) and annealed for 30 minutes at 37°C and 5 min 95°C. 1 μl phosphoannealed oligo (diluted 1:500) was then ligated into 1 μl of BsmBI-digested lentiviral pHK09-puro plasmid using 1 μl Quick ligase (NEB), 5 μl 2x Quick Ligase buffer (NEB) and 2 μl ddH2O for 5 minutes at room temperature. Guide RNA target sequences are provided in Table S2.

Pooled lentivirus was produced by co-transfecting equal amounts of each of these guide RNAs together with the psPAX2 and VSV.G vectors as described above. Virus was concentrated using a Beckmann XL-90 ultracentrifuge (Beckman Coulter) at 100,000 g (24,000 rpm) for 2 hours at 4°C. Viral titers were determined using alamarBlue staining, as described (https://portals.broadinstitute.org/gpp/public/resources/protocols). CCRF-CEM cells (40,000 per well in 100 µl RPMI medium) infected with lentiCas9-blast (a gift from Feng Zhang; addgene plasmid # 52962) were plated in 96-well format and transduced with lentivirus at multiplicity of infection (MOI) of 0.3. Infected cells were selected 48 hours post-infection with puromycin at 1 µg/ml for 7 days. Infected cells were treated with vehicle (PBS) or asparaginase in 24-well format (400,000 cell per well in 1 ml RPMI medium). Cells were harvested 5 days after start of asparaginase treatment, and genomic DNA was extracted using DNeasy Blood and Tissue Kit (Qiagen). Guide RNA sequences were PCR-amplified using pHKO9 sequencing primers (Table S2), PCR-purified using the QIAquick PCR purification Kit (Qiagen), and next-generation “CRISPR sequencing” was performed at the MGH CCIB DNA Core facility (https://dnacore.mgh.harvard.edu/new-cgi-bin/site/pages/crispr_sequencing_main.jsp). Cutting efficiency was assessed using CrispRVariantsLite v1.1 (Lindsay et al., 2016).

### Genome-Wide Loss of Function Screen

The genome-wide loss-of function screen was performed using the GeCKO v2 human library (a gift from Feng Zhang; Addgene #1000000048 and 1000000049), as described (Sanjana et al., 2014; Shalem et al., 2014). CCRF-CEM cells were first transduced with lentiCas9-blast, selected with blasticidin, and Cas9 activity was confirmed using a self-excising GFP construct, pXPR_ 011 (a gift from John Doench and David Root; addgene plasmid # 59702). GeCKO v2 consists of two half-libraries (A and B), each of which was transduced in biologic duplicates into 1.8 × 10^8^ Cas9-expressing CCRF-CEM cells at MOI = 0.3. Cells were selected with puromycin (1 µg/ml) beginning 24 hours post transduction, which was continued for 8 days. Based on the number of cells that survived selection and estimated growth rate, coverage was estimated at 663x for GeCKO half-library A, and 891x for GeCKO half-library B. Cells were split every other day, and the minimum number of cells kept at each split was 84 million for each half-library, in order to minimize loss of guide RNA coverage. Cells were treated with 10 U/L asparaginase beginning on day 10, and cells were harvested after 5 days of treatment. Genomic DNA was extracted using the Blood & Cell Culture DNA Maxi Kit (Qiagen). Samples were sequenced using CRISPR sequencing at the MGH CCIB DNA Core facility as described above. Significance of gene depletion based on guide RNA drop-out was calculated using MAGeCK software (https://sourceforge.net/projects/mageck/) (Li et al., 2014).

### shRNA and Expression Plasmids

The following lentiviral shRNA vectors in pLKO.1 with puromycin resistance were generated by the RNAi Consortium library of the Broad Institute, and obtained from Sigma-Aldrich:shLuciferase (TRCN0000072243), shNKD2#1 (TRCN0000187580), shNKD2#3 (TRCN0000428381), shLGR6#2 (TRCN0000063619) shLGR6#4 (TRCN0000063621), shAPC#1 (TRCN0000010296), shAPC#2 (TRCN0000010297), shGSK3α #1 (TRCN0000010340), shGSK3α #4 (TRCN0000038682), shGSK3α #2 (TRCN0000039564), shGSK3 β #6 (TRCN0000010551).

The construct encoding a constitutively active β-catenin (ΔN90) allele was a gift from Bob Weinberg (https://www.addgene.org/36985/). Expression constructs expressing wild-type FBXW7 (also known as CDC4) or its R465C mutant were a gift from Bert Vogelstein (https://www.addgene.org/16652/ and https://www.addgene.org/16653/). The 7TGC (TxTcf-eGFP//SV40-mCherry) TOPflash reporter was a gift from Roel Nusse (https://www.addgene.org/24304/). A hyperactive open-gate mutant of the human proteasomal subunit PSMA4 harboring a deletion of the N-terminal amino acids 2-10 (based on isoform NP_ 002780.1), termed ΔN-PSMA4, was designed based on the data of Choi and colleagues (Choi et al., 2016), and synthesized with a C-terminal V5 tag in a pLX304 lentiviral expression vector by GeneCopoeia (Rockville, MD). The GSK3α pWZL expression vector was previously described (Banerji et al., 2012).

### Assessment of Chemotherapy Response and Apoptosis

Cells (25,000 per well) were seeded in 100Δl of complete growth medium in 96-well plates and incubated with chemotherapeutic agents or vehicle. T-ALL cells were split every 48 hours in a 1:5 ratio. Briefly, 20 Δl of cells were mixed with 80 Δl fresh culture medium, supplemented with vehicle or chemotherapeutic drugs at the indicated doses. AML cells were split every 72 hours as described above. Cell viability was assessed by counting viable cells based on trypan blue vital dye staining (Invitrogen), according to the manufacturer’s instructions. Chemotherapeutic drugs included: Asparaginase (pegaspargase, Shire, Lexington, MA), dexamethasone (Sigma-Aldrich),vincristine (Selleckchem, Houston, TX), doxorubicin (Sigma-Aldrich), 6-mercaptopurine (Abcam, Cambridge, UK), CHIR99021 (Selleckchem), rapamycin (Selleckchem), RAD001 (Selleckchem), AZD2014 (Selleckchem), and Wnt3A (R&D systems, Minneapolis, MN). BRD0705 and BRD3731 were synthesized as described (Wagner et al., 2018). Caspase 3/7 activity was assessed using the Caspase Glo 3/7 Assay (Promega, Madison, WI) according to the manufacturer’s instructions.

### TOPFlash Reporter

Briefly, 400,000 CCRF-CEM cells were transduced with the TOPflash reporter 7TGC (Fuerer and Nusse, 2010), as described in the lentiviral infection section above. Cells expression the constitutive mCherry selection marker were selected by fluorescence activated cell sorting (FACS), treated with the Wnt ligand Wnt3A for 5 days, and cells with Wnt-inducible GFP expression were selected by sorting for mCherry and GFP double-positive cells. Selected cells were released from the Wnt ligand for a minimum of 7 days and then further manipulated with expression plasmids. GFP positivity of the mCherry positive cell fraction was assessed on a Beckton-Dickinson LSR-II instrument.

### Western Blot Analysis

Cells were lysed in RIPA buffer (Merck Millipore) supplemented with cOmplete protease inhibitor (Roche, Basel, Switzerland) and PhosSTOP phosphatase inhibitor (Roche). Laemmli sample buffer (Bio-Rad, Hercules, CA) and β-mercaptoethanol (Sigma-Aldrich) were mixed with 20 μg of protein lysate before being run on a 4–12% bis-tris polyacrylamide gel (Thermo Fisher Scientific). Blots were transferred to PVDF membrane (Thermo Fisher Scientific) and blocked with 5% BSA (New England Biolabs) in phosphate-buffered saline with 0.1% Tween (Boston Bioproducts, Ashland, MA) and probed with the following antibodies: Non-phospho β-catenin (Ser33/37/Thr41) antibody (1:1000, Cell Signaling Technologies #8814), total β-catenin antibody (1:1000, Cell Signaling Technologies #8480), total GSK3α/β antibody (1:1000, Cell Signaling Technologies #5676), phospho-GSK3α/β (Tyr279/216) antibody (Thermo Fisher Scientific #OPA1-03083), Myc-tag antibody (1:1000, Cell Signaling Technologies #2272), phospho-p70 S6 kinase (Thr389) antibody (1:1000 Cell Signaling Technologies #9234) or GAPDH (1:1000, Cell Signaling Technologies #2118). Detection of horseradish peroxidase-linked secondary antibodies (1:2000, Cell Signaling Technologies #7074S) with horseradish peroxidase substrate (Thermo Fisher Scientific) was visualized using Amersham Imager 600 (GE Healthcare Life Sciences, Marlborough, MA).

### Assessment of Cell Size

Cells (400,000 per well) were plated in 1ml of complete growth medium, containing a final concentration of 10 U/L asparaginase in a 24-well format. After 48 hours of treatment, forward scattering (FSC-H) was assessed by flow cytometry.

### Quantitative Reverse Transcriptase PCR (qRT-PCR)

RNA was isolated using RNeasy kit (Qiagen) and cDNA was made using SuperScript III first-strand cDNA synthesis kit (Thermo Fisher Scientific). qRT-PCR was performed using Power SYBR® Green PCR Master Mix (Thermo Fisher Scientific) and 7500 real-time PCR system (Applied Biosystems). Primers used are described in Table S2.

### *In Vivo* Drug Treatment of Patient-Derived Xenograft

T-ALL clinical specimens were collected from children with newly diagnosed T-ALL enrolled on Dana-Farber Cancer Institute clinical trials, with informed consent and institutional review board approval in accordance with the Declaration of Helsinki. Patient-derived xenografts were generated by engraftment viably frozen leukemic cells into immunodeficient mice, as described (Townsend et al., 2016).

NRG mice were exposed to a sub-lethal dose of radiation (4.5 Gy) and subsequently injected with human leukemia cells. All injections were performed by tail vein injection. Blood collections were performed to monitor leukemia onset in mice injected with leukemic cells, which was assessed by staining for human CD45 expression using anti-human CD45-PE-Cy7 (1:400, BD# 560915) antibody staining. As soon as leukemia engraftment was confirmed based on ≥5% human leukemia cells in the peripheral blood, treatment of mice was begun. Asparaginase (1,000 U/kg) or PBS were injected by tail vein injection as a single dose on day 1 of drug treatment, and BRD0705 (15 mg/kg) or vehicle were given every 12 hours for 12 days by oral gavage. Vehicle was formulated as previously described (Wagner et al., 2018). Mice were anesthetized with isoflurane (Patterson Veterinary, Greeley, CO) prior to gavage. After start of treatment, body weight was monitored every third day. To assess bilirubin levels, mandibular blood vein collection was performed, and bilirubin levels were measured in the Boston Children’s Hospital clinical laboratory. The primary endpoint of this experiment was survival. Mice were euthanized as soon as they developed weight loss greater than 15% or signs/symptoms of progressing disease. Post-mortem analysis of leukemic burden was performed by extracting bone marrow cells, which were filtered through a 40 μM mesh filter and red blood cells were lysed using the BD Biosciences Red Blood Cell Lysis reagent (BD #555899). Isolated bone marrow cells were stained for assessment of leukemic burden using following antibodies from BD Biosciences: Anti-human CD4-APC-Cy7 (1:100, BD# 561839) and anti-human CD8-PerCP-Cy5.5 (1:100, BD# 560662).

### Assessment of Protein Stability

Protein degradation was assessed using a non-radioactive quantification of the methionine analog L-azidohomoalanine (AHA) AlexaFluor488 (Thermo Fisher), as previously described (Wang et al., 2017). Briefly, 400,000 CCRF-CEM cells were seeded in 1 ml of methionine-free RPMI containing 10% dialyzed FBS. After 30 minutes, the pulse step was performed by replacing this media with 1 ml RPMI supplemented with 10% dialyzed FBS and AHA at a final concentration of 50 µM for 18 hours. In the chase step, cells were released from AHA by replacing media with RPMI containing 10% dialyzed FBS and 10x L-methionine (2 mM) for 2 hours. Subsequently, media was replaced with regular growth medium and cells were treated with a final concentration of 10 U/L asparaginase, followed by fixation of cells. AHA labeled proteins were tagged using TAMRA alkyne click chemistry, and fluorescence intensity was measured by flow cytometry. A sample without AHA labeling but TAMRA alkyne tag was included as a negative control to account for background fluorescence.

## QUANTIFICATION AND STATISTICAL ANALYSIS

For two-group comparisons of continuous measures, a two-tailed Welch unequal variances t-test was used. For 3-group comparisons, a one-way analysis of variance model (ANOVA) was performed and a Dunnett adjustment for multiple comparisons was used. For analysis of two effects, a two-way ANOVA model was constructed and unless indicated included an interaction term between the two effects. Post-hoc adjustment for multiple comparisons for two-way ANOVA included Tukey’s adjustment. The log rank test was used to test for differences in survival between groups, and the method of Kaplan and Meier was used to construct survival curves. Data shown as bar graphs represent the mean and standard error of the mean (s.e.m) of a minimum of 3 biologic replicates. All p-values reported are two-sided and considered as significant if <0.05.

